## Supplementary Materials for "Repeated MRI scans of the human brain: diurnal oscillations in healthy adults and bipolar disorder patients"

### **Supplementary Information**

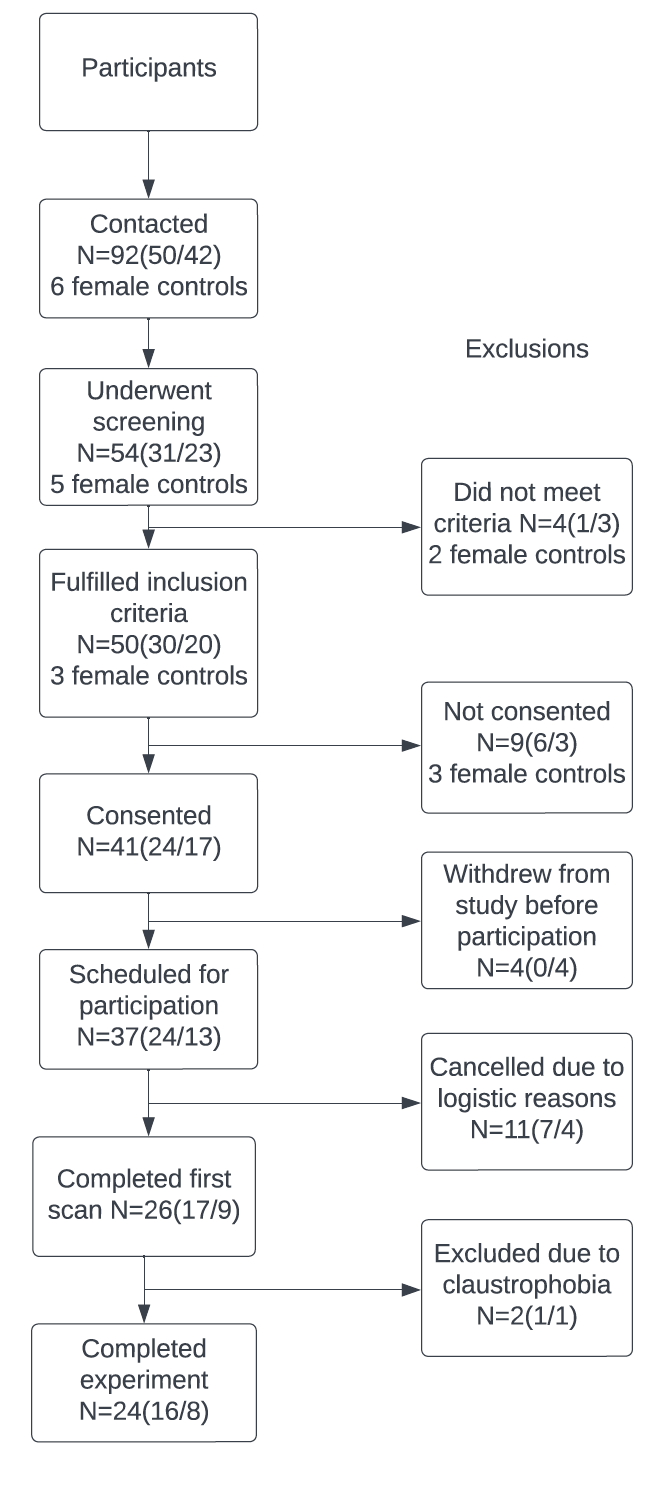

**Supplementary Fig. 1 |** **Recruitment and retention flowchart.**

Numbers (n/n) in the parentheses are controls (first) and patients with BPD (second).

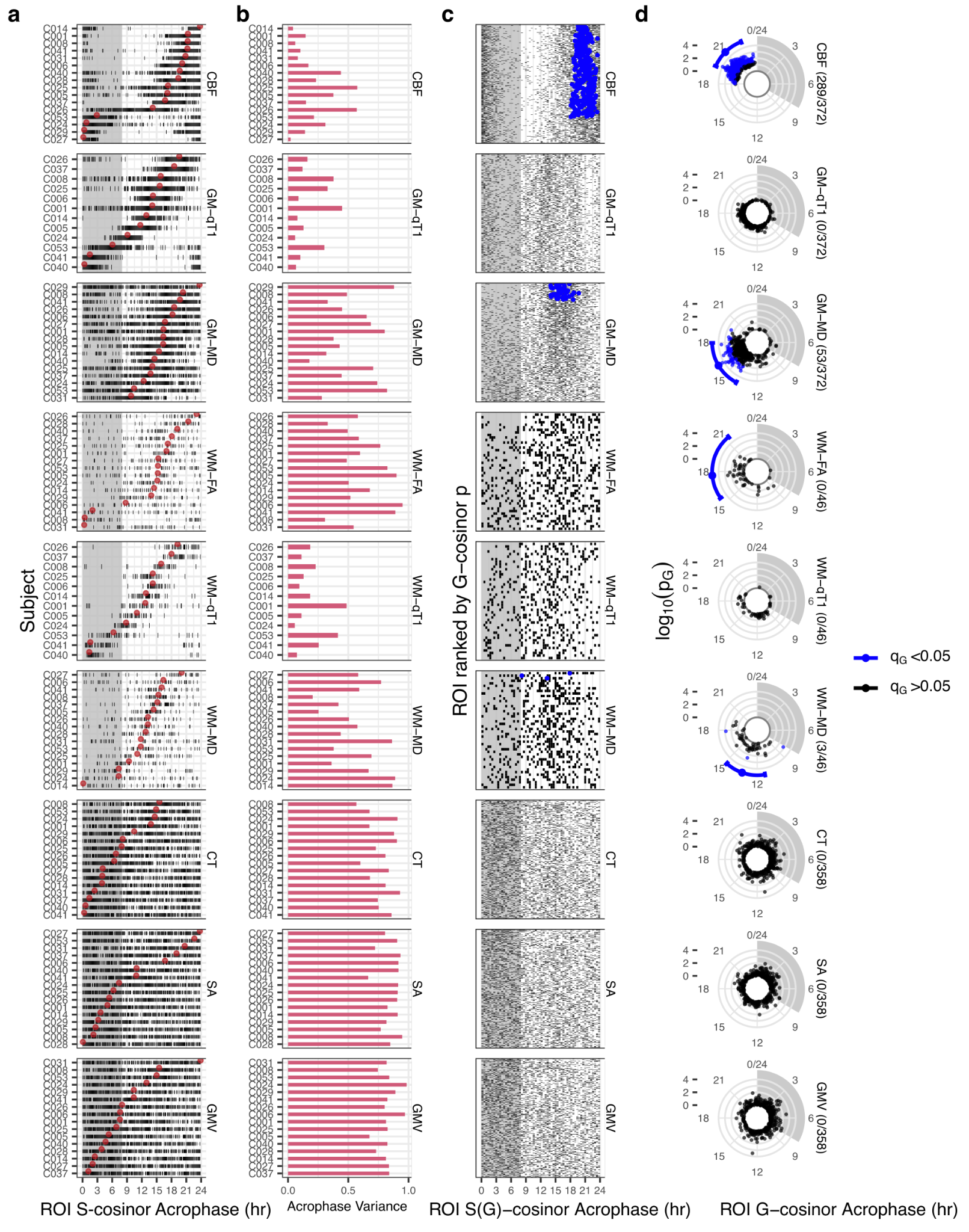

**Supplementary Fig. 2 | Regional cosinor statistics for all metrics.**

**a-b)** As described in Figure 3a. **c)** For each ROI (y-axis - sorted by ROI G-cosinor p-value within each metric), each subject’s cosinor acrophase estimate (x-axis) is shown as a black tick to illustrate within-ROI consistency. ROIs with G-cosinor significant acrophases (q<0.05) are shown as blue dots. **d)** As described in Figure 3c.

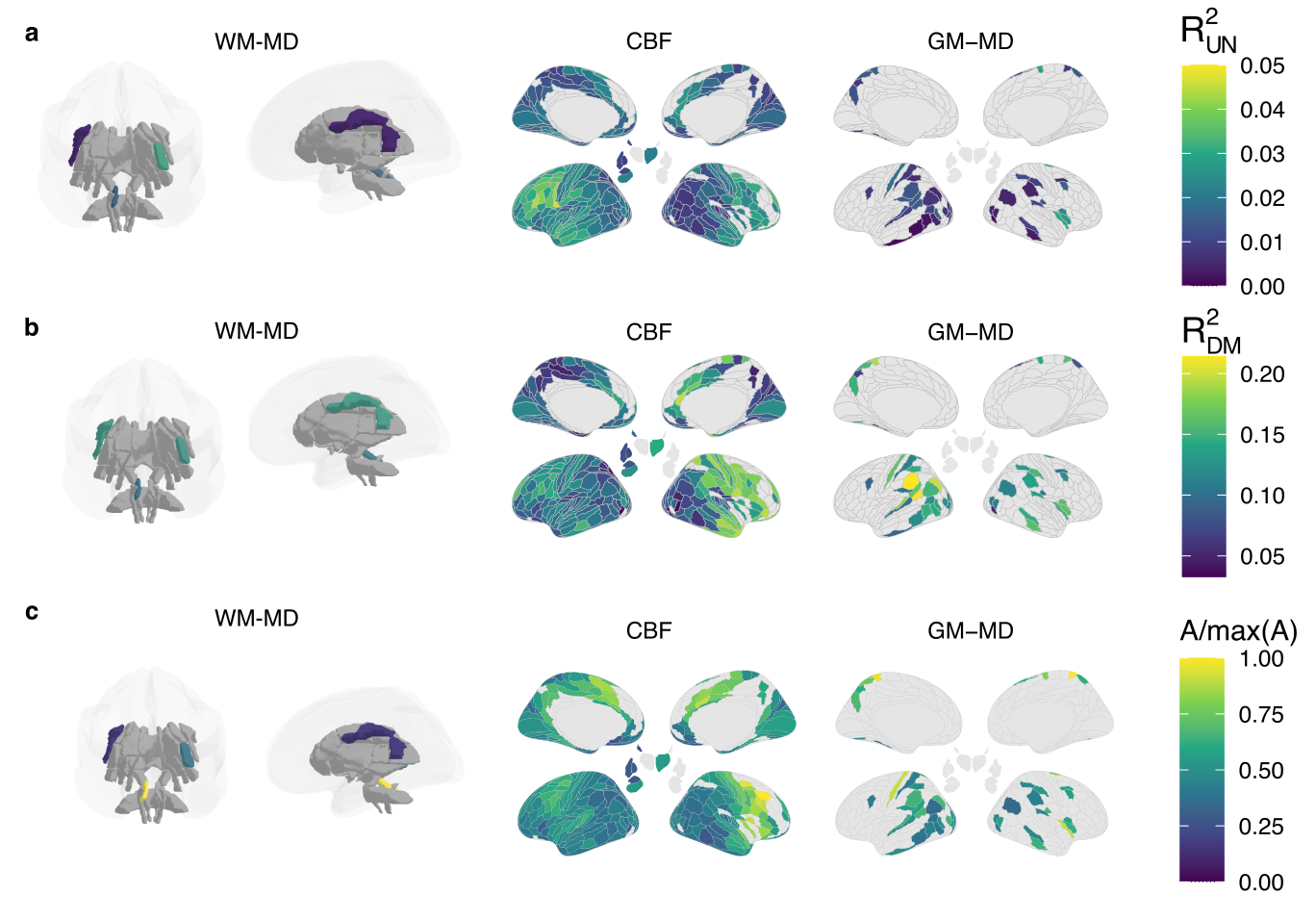

**Supplementary Fig. 3 | Regional G-cosinor statistics in controls.**

**a-c)** Spatial distribution of G-cosinor statistics for CBF, WM- and GM-MD regions surviving FDR-correction, q<0.05. **a)** R^2^_UN_, **b)** R^2^_DM_ and **c)** Amplitudes (A), scaled relative to the maximum amplitude within metric for visualization purposes.

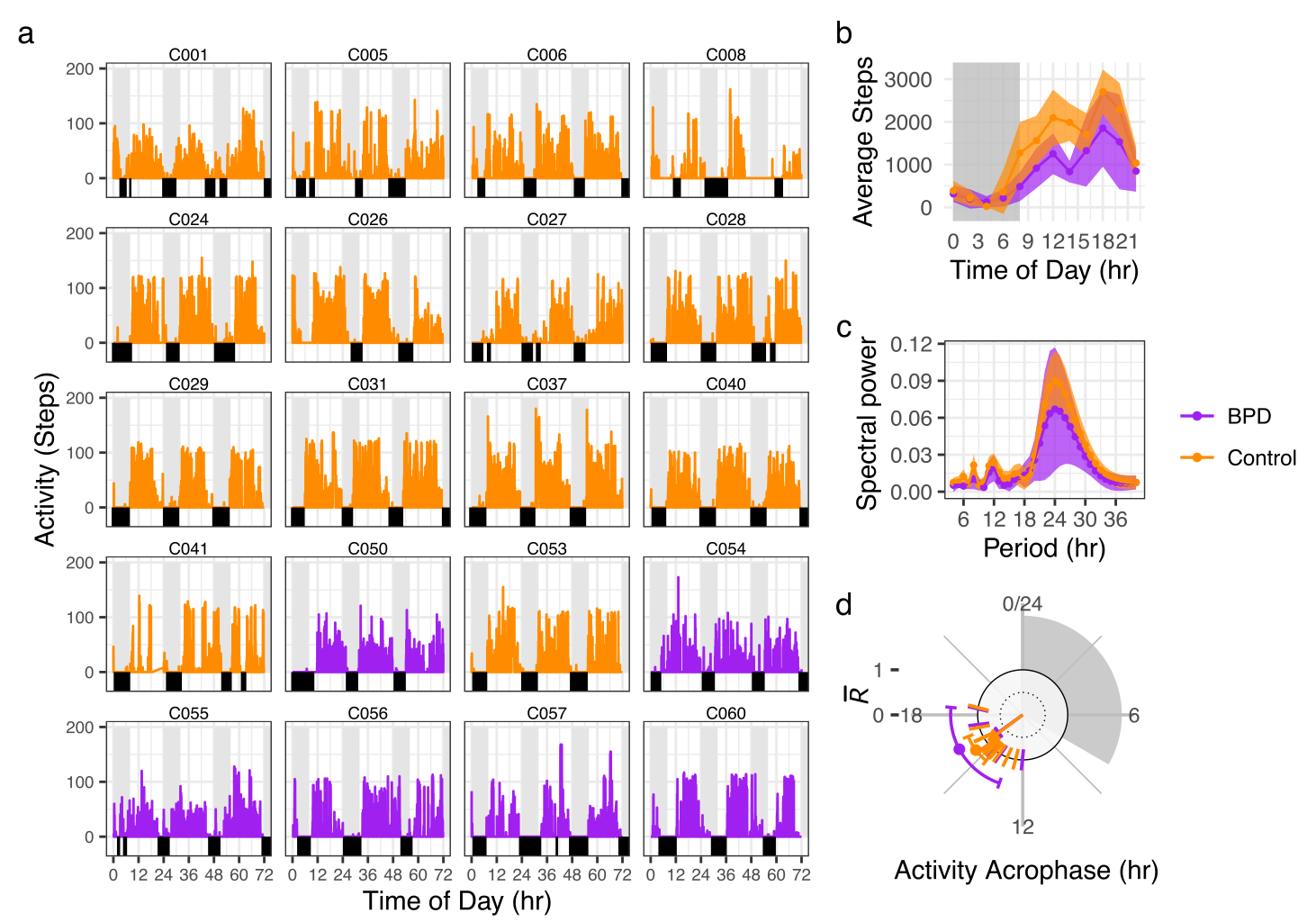
**Supplementary Fig. 4 | Actigraphy in units of steps registered.**

**a)** 3 days of raw step data from actigraphy with sleep (black bars). **b)** 2 hour averages with mean and 95% CI of the mean (confidence band). **c)** Spectral power of periods (R^2^ of S-cosinor fit) from 4hr to 40hr on step data for each subject averaged within group with mean and 95% CI of the mean (confidence band). **d)** Methods from Figure 4a applied to 1hr epoch actigraphy step data.

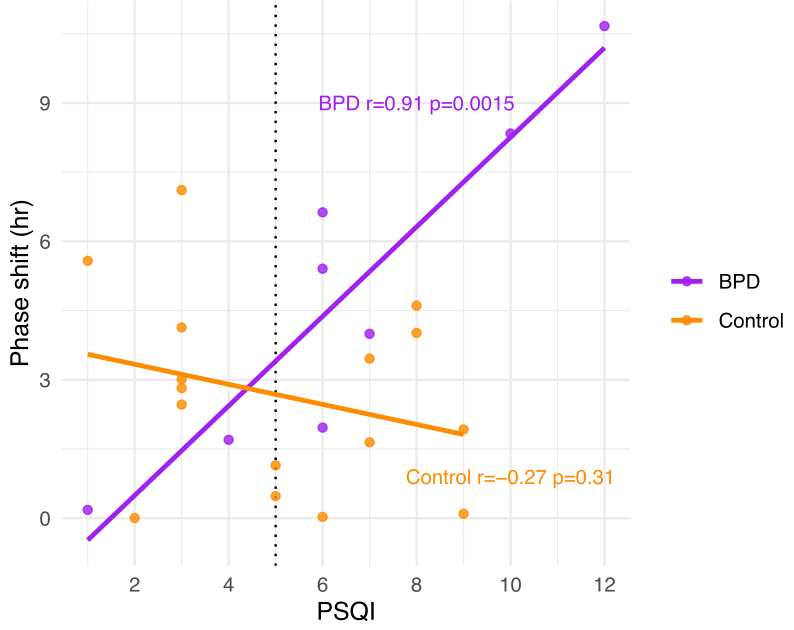

**Supplementary Fig. 5 | CBF phase deviation correlates with subjective sleep quality in BPD.**

Phase shift was calculated for each subject as the minimum arc length relative to the G-cosinor acrophase in the combined group. Sleep quality was measured with the PSQI, where higher values indicate poor sleep quality. The dotted vertical line indicates the PSQI threshold for good (<5) and poor (>5) sleep quality^1^. Each group was tested with Pearson’s correlation (results in panel).

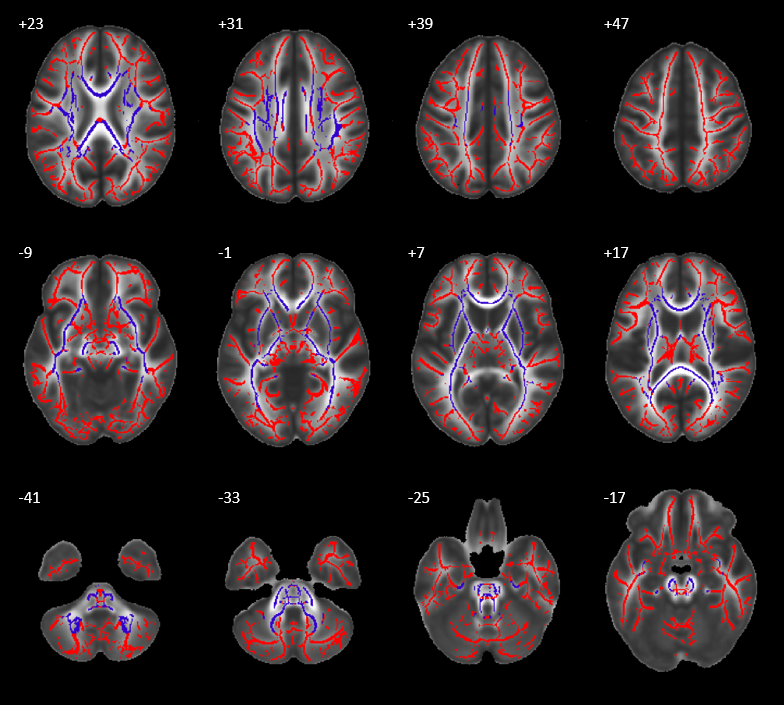

**Supplementary Fig. 6 | TBSS skeleton overlaid on the mean FA map.**

Blue: voxels included in the regional analyses, obtained from the JHU Atlas^2^. Red: additional voxels included in the whole-skeleton analysis. Slice locations are indicated at the upper left of each slice, left is on the left, and up is anterior.

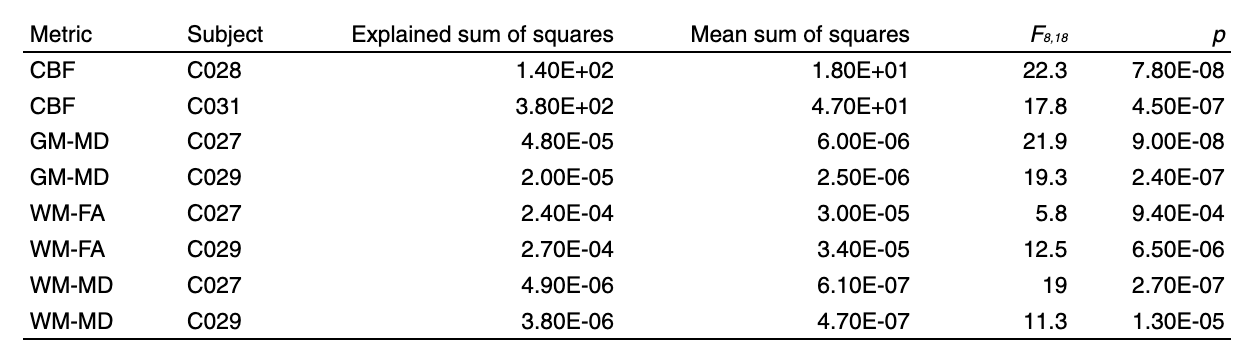
**Supplementary Table 1 | ANOVA on whole-brain-averaged back-to-back repeated scan data.** Diffusion tensor imaging or arterial spin labeling measurements were repeated three times at each of the nine time points in two subjects. A one-way ANOVA on whole-brain data assessing within- vs. between-time point variation was performed to determine if scan-to-scan variation was smaller than across-time variation.

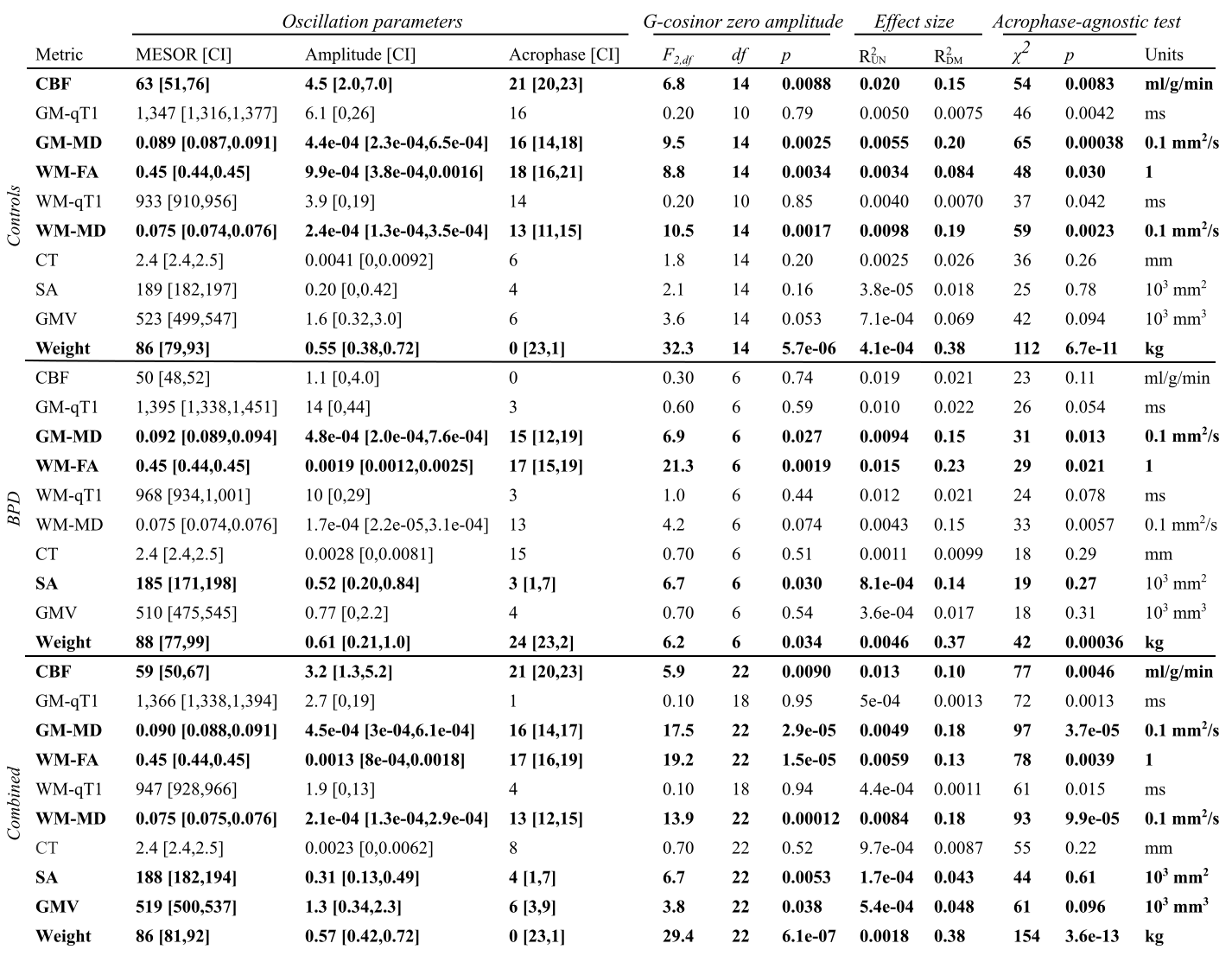

**Supplementary Table 2 | Cosinor statistics for all MRI metrics at the whole-brain level.**

When p >0.05, lower bounds of amplitude CI are shown as “0” and acrophase CI is not shown. Rows with G-cosinor p<0.05 are in bold.

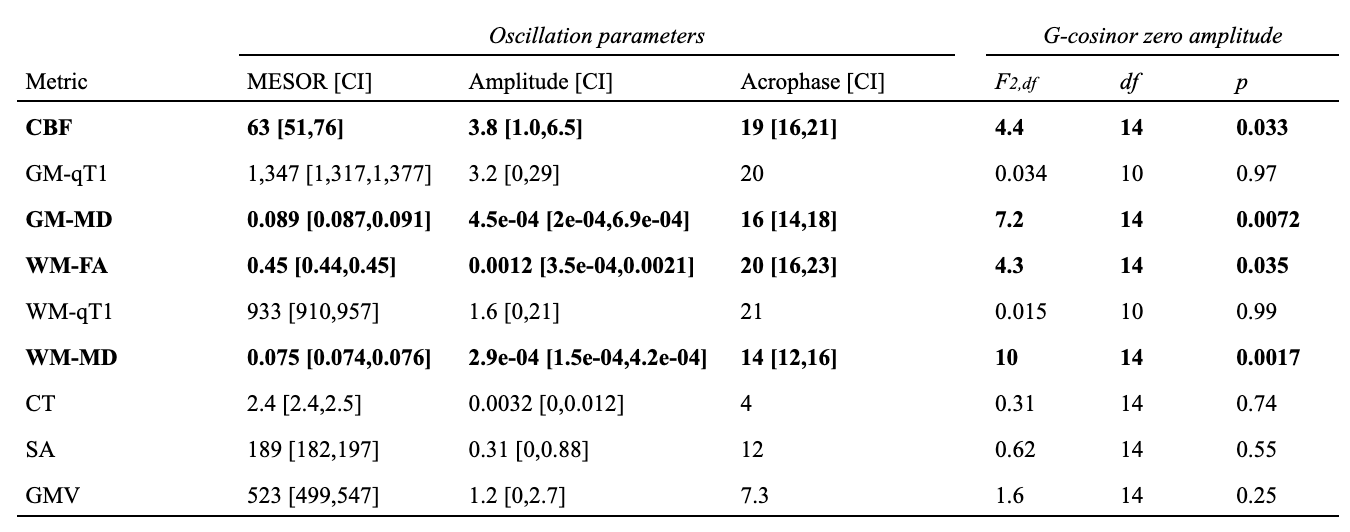

**Supplementary Table 3 | G-cosinor models after covarying for body weight**.

Each subject first had an S-cosinor model fit with their weight prior to each scanning session as an additional covariate. Amplitudes and acrophases of these fits were used for the G-cosinor F-test. Rows with G-cosinor p<0.05 are in bold.

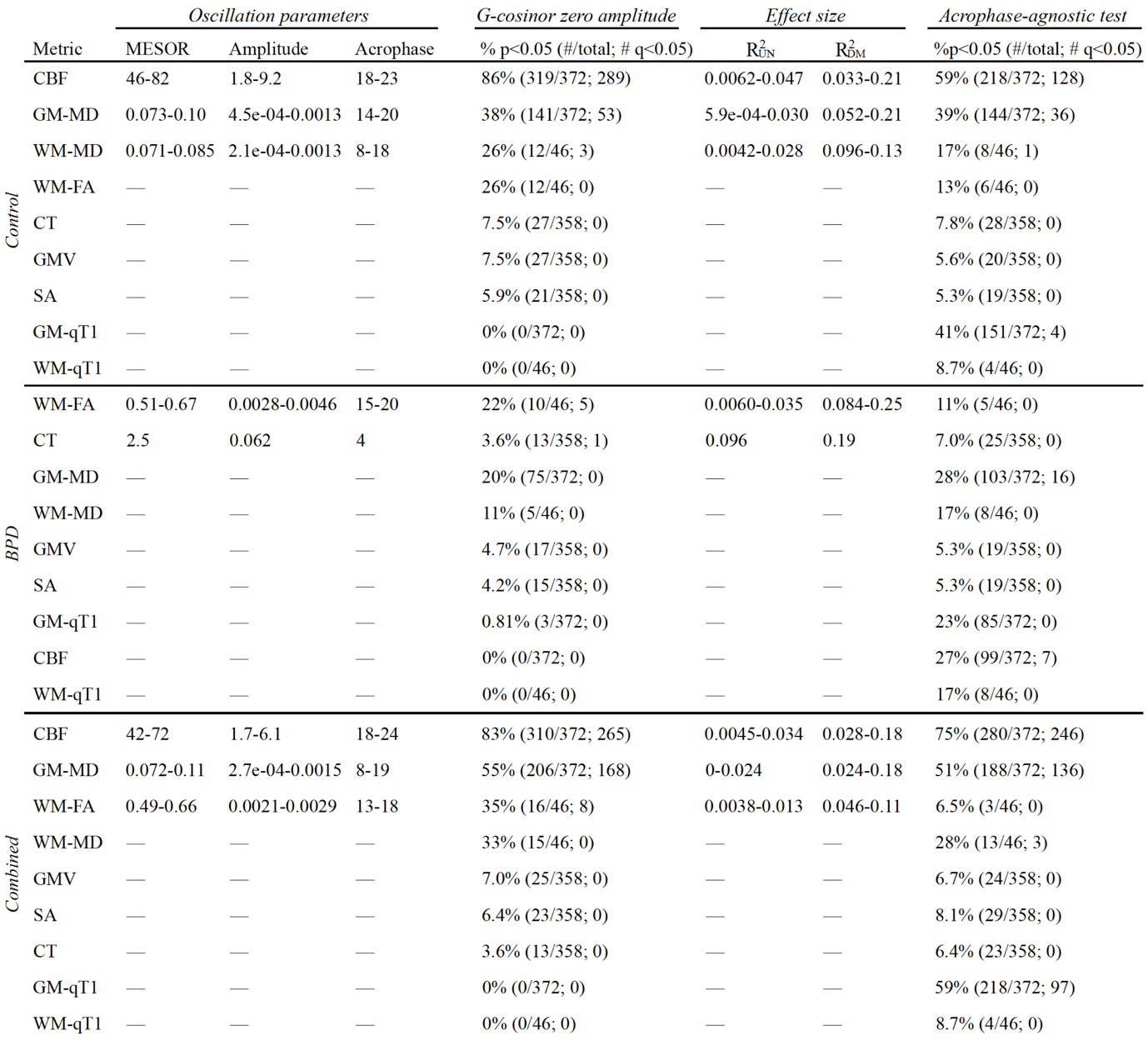
**Supplementary Table 4** | **Summary of cosinor statistics for all MRI metrics at the regional level.** Ranges are given for all G-cosinor FDR q<0.05 ROIs within each metric.
